# Intracellular production of hydrogels and synthetic RNA granules by multivalent enhancers

**DOI:** 10.1101/117572

**Authors:** Hideki Nakamura, Albert A. Lee, Ali Sobhi Afshar, Shigeki Watanabe, Elmer Rho, Shiva Razavi, Allison Suarez, Yu-Chun Lin, Makoto Tanigawa, Brian Huang, Robert DeRose, Diana Bobb, William Hong, Sandra B. Gabelli, John Goutsias, Takanari Inoue

**Author notes:** These authors contributed equally (Hideki Nakamura, Albert A. Lee). To whom general Correspondence should be addressed (T.I.). To whom correspondence regarding the computational analysis should be addressed (A.S.A).

## Abstract

Non-membrane bound, hydrogel-like entities, such as RNA granules, nucleate essential cellular functions through their unique physico-chemical properties. However, these intracellular hydrogels have not been as extensively studied as their extracellular counterparts, primarily due to technical challenges in probing these materials *in situ.* Here, by taking advantage of a chemically inducible dimerization paradigm, we developed iPOLYMER, a strategy for rapid induction of protein-based hydrogels inside living cells. A series of biochemical and biophysical characterizations, in conjunction with computational modeling, revealed that the polymer network formed in the cytosol resembles a physiological hydrogel-like entity that behaves as a size-dependent molecular sieve. We studied several properties of the gel and functionalized it with RNA binding motifs that sequester polyadenine-containing nucleotides to synthetically mimic RNA granules. Therefore, we here demonstrate that iPOLYMER presents a unique and powerful approach to synthetically reconstitute hydrogel-like structures including RNA granules in intact cells.

## Introduction

A hydrogel is a hydrophilic polymer network that is capable of absorbing water ^1^. An outstanding feature of hydrogels as a material is that their physico-chemical characteristics can be feasibly tuned over a wide range, by changing relevant parameters such as concentrations of polymers and cross- linkers, or ambient environments including temperature and pH. In a biological context, these highly variable properties enable biological hydrogel-like structures to serve versatile roles in living organisms, such as supporting and regulating functional entities ^2,3^ as well as lubricating joints ^4^. Recent works have found that biological hydrogel-like structures not only exist in extracellular space, but also inside cells ^5–7^. These intracellular hydrogels serve vital functions, such as forming diffusion barriers at the interface of subcellular compartments or nucleating cellular activities ^8–10^. Cytoskeletal filamentous networks including actin and intermediate filaments such as neurofilaments have also been implicated to form hydrogel-like structures, which are dynamically regulated to perform biologically essential processes such as cell migration ^11^, adhesion ^11^ and modulation of axonal transport ^12^.

A significant intracellular structure that has been related to hydrogels is the RNA granule. RNA granules are known to exhibit a phase separation-like behavior and be subject to dynamic structural rearrangements ^6,9,13,14^. Moreover, many of their components contain low complexity sequences, which actually form hydrogels when purified at higher concentrations ^6^. While RNA granules are physiologically important ^15–17^, their structural organization and biological relevance remain uncharacterized. Development of synthetic equivalents of the granules may become an alternative for facilitating our understanding of the relationship between the structure and function of these intracellular hydrogel-like structures.

Synthetic hydrogels have long been of great interest in the field of biomedical engineering, primarily because their physical properties can be designed to achieve a desired objective; e.g., to produce synthetic biomaterials that can become a surrogate for damaged tissue ^18,19^. Innovations in polymer and protein science have already enabled the development of numerous synthetic hydrogels that are successfully controlled in a stimuli-responsive manner ^20–23^ and are currently used in clinical practice and bioengineering research ^24,25^. However, past research has principally focused on extracellular applications, including the design of tissue engineering scaffolds and drug delivery vehicles ^25,26^. Until now, little has been achieved in an effort to generate synthetic hydrogels inside cells, mainly due to the challenging nature of inducing gel formation in intact living cells. As such, researchers currently resort to either microinjection of acrylamide gels already formed outside cells ^27^, or overexpression of building block molecules whose polymerization cannot be triggered ^28,29^. These reports employed the hydrogels as mechanical probes, described their dynamics in living cells, or evaluated the effects of gel formation on cell survival. Unfortunately, they have been less successful in directly demonstrating that synthetic hydrogels are biologically functional within living cells, primarily due to the lack of an experimental paradigm that is capable of forming gels in an inducible manner, with sufficiently fast kinetics to allow monitoring functionality before and after induction.

Another challenge in studying hydrogel formation inside cells is the limited toolkits available for gel evaluation. It is often not straightforward to claim the identity of an object observed in cells as being a gel, owing to limited physical access. Reconstitution of the material *in vitro* is the most frequently adopted way to address this issue ^6,8,14,30^ although the conditions adopted *in vitro* may not necessarily recapitulate the phenomena observed in living cells. Comprehensive understanding of the nature of induced hydrogels therefore requires the development of a totally new paradigm.

## Results

### Design principle of iPOLYMER strategy

To enable hydrogel formation inside living cells in a rapidly inducible manner without committing to invasive approaches such as microinjection, we introduce a novel strategy, termed iPOLYMER, for <u>i</u>ntracellular <u>p</u>roduction <u>o</u>f <u>l</u>igand-<u>y</u>ielded <u>m</u>ultivalent <u>e</u>nhancers. By definition, a hydrogel is formed by polymers held together either via cross-linkers or via direct physical or chemical interactions between them. We thus utilized a chemically inducible dimerization (CID) technique ^31^ to cross-link two molecular species interlaced with peptide chains. In this system, the chemical agent rapamycin induces dimerization between two proteins, the FK506 binding protein (FKBP) and the FKBP-rapamycin binding protein (FRB), with high specificity and fast kinetics (Fig. 1a). Our group and others have exploited CID and confirmed its robustness in protein dimerization as well as its versatility in achieving rapid dimerization of proteins of interest in living cells ^31^. The CID may thus offer an ideal building block for inducing *in situ* gel formation. We hypothesized that multiple copies of FKBP and FRB molecules, interspaced with a polypeptide chain, should undergo polymerization when expressed in cells and exposed to rapamycin, leading to a hydrogel-like network induced inside living cells (Fig. 1b).

**Figure 1.**
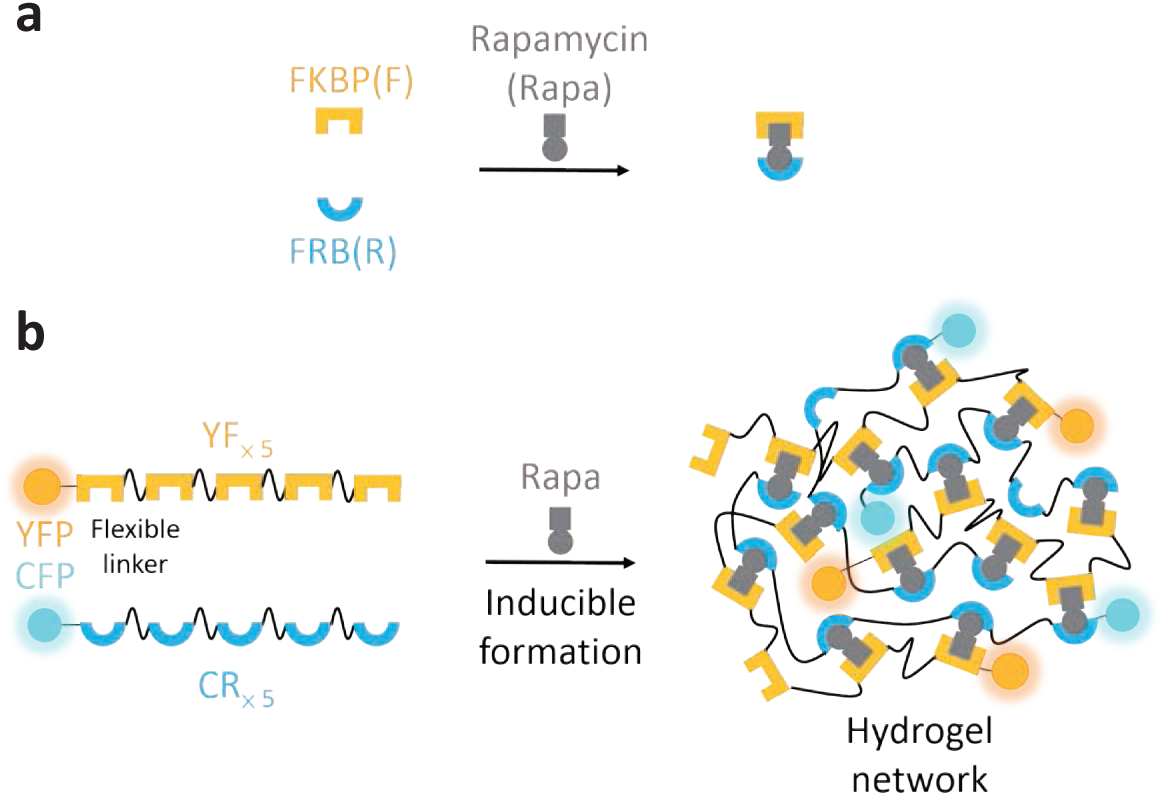
Schematic illustration of iPOLYMER. **(a)** Rapamycin induces rapid, stable and specific binding between FKBP and FRB molecules. **(b)** Two series of proteins, YF_xN_ and CR_xM_, were engineered to track their expression in cells: a yellow fluorescent protein (YFP) on up to five tandem repeats of an FKBP domain and a cyan fluorescent protein (CFP) on up to five tandem repeats on an FRB, spaced by 12 amino acid linker sequences. Mixing YF_x5_ and CR_x5_ (left) with rapamycin is expected to induce the formation of a hydrogel network (right). YF_XN_ and CR_XM_ contain N-repeats of FKBP and M-repeats of FRB with the same linkers, respectively.

### Computational model reveals size distribution and valence number dependency of polymer network

We first explored the feasibility of iPOLYMER for rapid hydrogel-like network synthesis *in silico* by carrying out kinetic Monte Carlo simulations based on a realistic stochastic physical model of three- component multivalent-multivalent molecular interactions (Fig. 2a), whose association/dissociation rates were determined from previous experimental findings ^32^ (see Materials and Methods and Supplementary Methods for details). Simulation results for different valencies of equivalent FKBP and FRB proteins demonstrated, for higher valencies of three or more, quick formation of relatively large aggregates comprising many FKBP and FRB molecules occurred, while only small aggregates were seen in valence number two, and none observed for valence number one, as expected (Supplementary Movie 1-5). Moreover, convergence to a stationary state was observed, characterized by the formation of a single hydrogel-like aggregate that may coexist with much smaller molecules.

**Figure 2.**
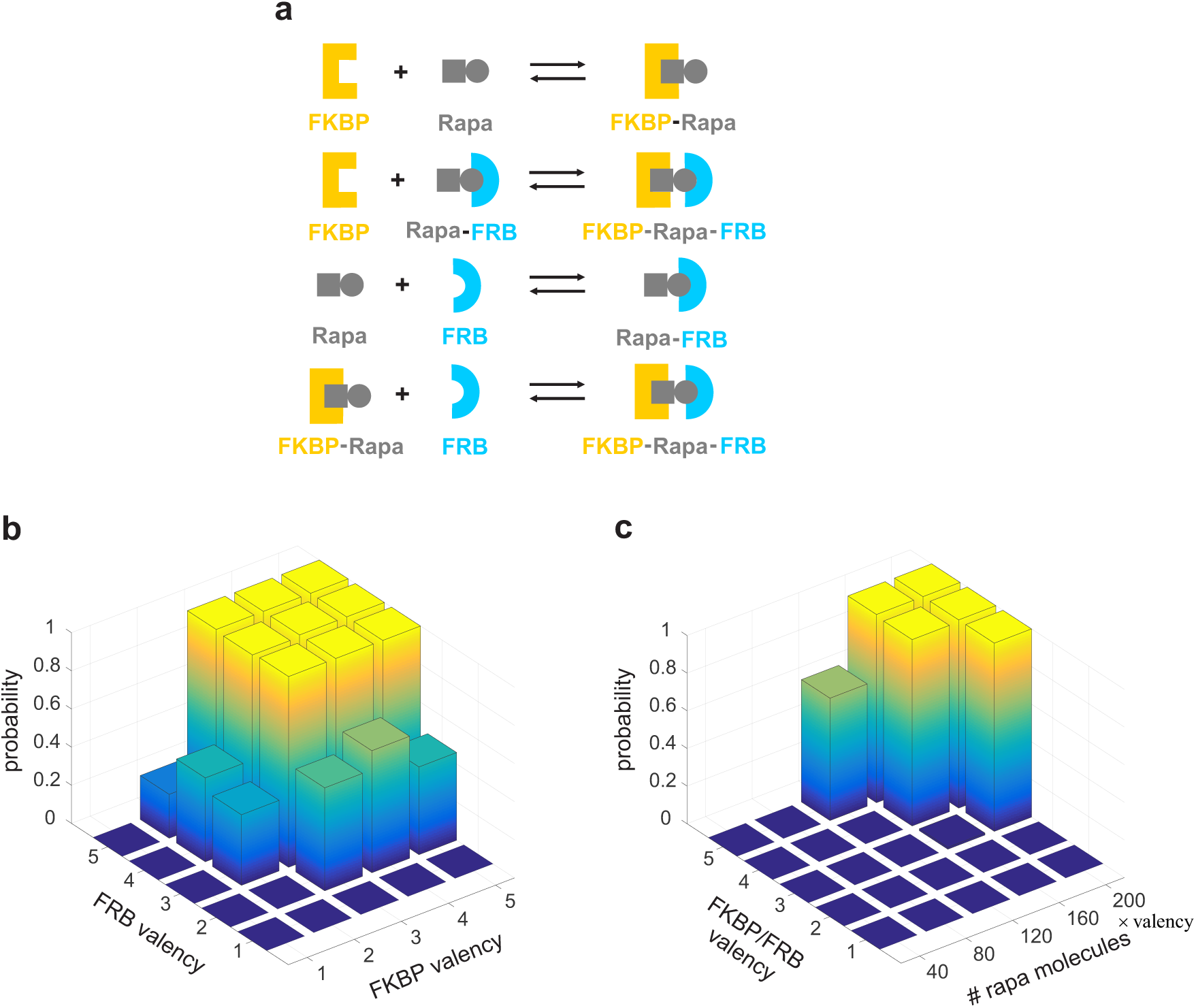
*In silico* implementation of iPOLYMER demonstrates its feasibility for hydrogel network synthesis. (a) Four reversible reactions between monomeric FKBP, FRB and rapamycin molecules modeled in our simulations. Each binding unit in the tandem repeats of FKBP or FRB can undergo the four reactions in the presence of rapamycin. (b) Estimated probabilities that iPOLYMER will produce aggregates of a threshold size of 100 or larger for different valence numbers of the FKBP and FRB molecules. An aggregate of size 100, as defined in the Supplementary Methods, comprises 25% of the total number of FKBP and FRB molecules initially present in the simulated system. The sharp increase in the probability values indicates that efficient polymerization can be achieved when the individual valence numbers of FKBP and FRB are at least three, with the total valence number of FKBP and FRB molecules being at least six. (c) Estimated probabilities that iPOLYMER will produce aggregates of a threshold size of 100 or larger for different valence numbers of the FKBP and the FRB molecules and different numbers of rapamycin molecules, determined for each simulation by multiplying a base number of rapamycin molecules with the common valency of FKBP and FRB [e.g., base rapa # (160) x valency (4) = rapa # (640)] in order to scale the effect of peptide valency on the number of binding sites against rapamycin. The observed sharp decrease in the probability values indicates that efficient polymerization requires a sufficient concentration of rapamycin. This implies that, in addition to the valence numbers of FKBP and FRB, the concentration of the dimerizing agent is expected to directly affect phase transition.

We then computed the size distribution of the molecular aggregates at six equally-spaced time points within a predefined time interval (Supplementary Methods, Figs. SM3-SM7). These distributions effectively summarize the formation of aggregates from smaller molecules and demonstrate the fact that iPOLYMER can gradually yield aggregate molecules. At valence numbers three or more, the size distribution dramatically spreads over larger sizes, reflecting the formation of increasingly large aggregates. As the valence number increases, more aggregates with larger sizes are gradually formed, highlighting the crucial role that valency plays in the formation of complex aggregates.

The observed strong dependency of aggregate formation on valency was consistent with previous theoretical and experimental findings, which have implied that a sharp phase transition from small assemblies to macroscopic polymer gels takes place when the degree of bonding increases ^28,33^. By simulating the process of aggregate formation using our realistic physical model, we further demonstrated that, at valence number three or more, the initial unimodal size distribution transitions into a bimodal distribution, indicating the coexistence of two distinct phases (Supplementary Methods, Figs. SM5-SM7). Further evidence of phase transition was also demonstrated by the estimated probabilities of iPOLYMER to produce aggregates of a threshold size of 100 or larger for different valence numbers (Fig. 2b), as well as for different rapamycin concentrations (Fig. 2c).

### *In silico* analysis predicts molecular sieving property with estimated pore sizes

Molecular aggregates are often known to form a selective three-dimensional gel-like sieve ^3,34^ To investigate the sieving properties of molecular aggregates produced by iPOLYMER *in silico,* we constructed graph representations of such aggregates (Supplementary Methods, Figs. SM8 and SM9), and used these representations to define the *effective pore size* (EPS) of a sieve as a measure for quantifying the sizes of molecules that will most likely pass through the sieve. Allthough we could potentially derive this measure from the *pore size distribution* (PSD) of a molecular aggregate (Supplementary Methods), calculating PSDs for non-trivial dense aggregates, which are the types of aggregates observed at steady state in CID systems in which phase transition takes place, is a computationally intractable problem. To tackle this issue, and consistent with our experiments, we first considered a CID system comprising a sufficient number of rapamycin molecules, as well as tandem FKBP and FRB peptides of valence 5. We then utilized experimentally determined lengths of the peptides forming an aggregate and computationally inferred a range of plausible EPS values (10-28 nm) for such aggregates by taking into account the dense structure of the graphs associated with molecular aggregates observed at steady state (Supplementary Methods).

Because of difficulties in experimentally estimating the EPS value of a molecular aggregate at an early stage of its formation, we sought to derive this value by approximately computing the associated PSD, in order to gain further insight into the aggregate's properties (Supplementary Methods). *In silico* analysis revealed that the PSD and EPS of an early-stage aggregate is directly influenced by the valence number of the FKBP and FRB molecules. We specifically demonstrated that polymerization of FKBP and FRB molecules with larger valences result in early-stage aggregates with coarser sieving potential than molecular sieves formed by molecules with smaller valences (Supplementary Methods, Figs. SM11 and SM12), although this trend was argued to be reversed at steady state, provided that a sufficient concentration of rapamycin molecules was present.

### iPOLYMER aggregate formed in living cells

To test our computational results, we performed experiments to evaluate iPOLYMER in living cells. Towards this end, we generated two series of engineered proteins in order to track their expression in cells (Fig. 1b): a yellow fluorescent protein (YFP), on up to five tandem repeats of an FKBP domain (YF_xN_, N = 1,2,3,4,5), and a cyan fluorescent protein (CFP) on similar tandem repeats of an FRB domain (CR_xM_, M = 1,2,3,4,5). We co-expressed in COS-7 cells the highest-valence number pair, YF_x5_ and CR_x5_, confirmed their distribution in the cytosol, and added rapamycin while performing live-cell fluorescence imaging. In cells with high expression of both peptides, YF_x5_ and CR_x5_ initially exhibited diffuse fluorescence signals that rapidly turned into puncta upon rapamycin addition (Fig. 3a). These puncta steadily grew in size during prolonged rapamycin treatment. We carried out the experiment with a Förster resonance energy transfer (FRET) measurement between CFP and YFP on the proteins and observed increasing FRET values in the cytosol within 5 min of rapamycin addition, whereas a continuous FRET increase was observed at the puncta that emerged at later time points (Fig. 3a, Supplementary Fig. 1, and Supplementary Video 6). This implied that puncta formation was due to the binding between the two proteins induced by rapamycin, which was further supported by the lack of puncta in DMSO-treated cells (Fig. 3a). FRET ratio fold-change during puncta formation was significantly greater at the puncta compared to that in the cytosol, while the fold-change in the cytosol was significant compared to that in DMSO control cells (Fig. 3a). We then examined if the growth of each punctum involved coalescence between multiple puncta, and found that two puncta occasionally coalesced into a larger one upon collision (Supplementary Fig. 2, upper panel). However, these puncta exhibited minimal reorganization of their apparent shape, reflecting the practically irreversible nature of rapamycin-induced CID in the cell ^35^. Due to continuous coalescence of the aggregate, we observed large aggregates typically surrounding nuclei in the cell treated with rapamycin for 24 h (Supplementary Fig. 2, lower panel).

**Figure 3.**
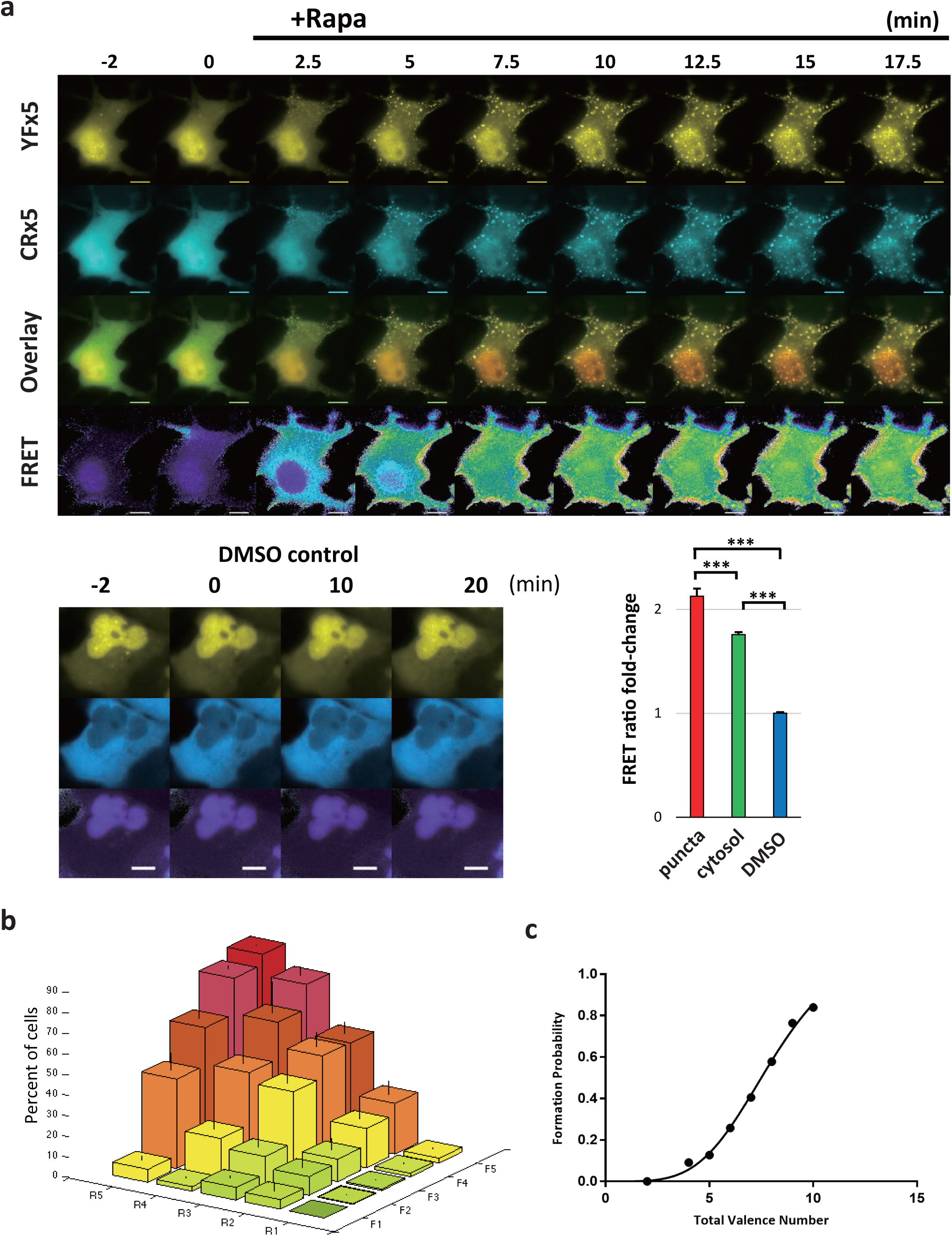
iPOLYMER puncta formation in living cells. **(a)** Time-lapse imaging of fluorescent puncta formation in COS-7 cells at indicated times relative to the addition of rapamycin. Scale bars: 10 μm. Punctate structures enriched with CFP, YFP, and FRET signals start to emerge within 5 min after 333 nM rapamycin addition, while DMSO treated cells demonstrate lack of puncta formation. The FRET ratio fold-change was significantly greater at puncta compared to that in the cytosol, which in turn was significantly greater than in DMSO treated cells. **(b)** Frequency of iPOLYMER puncta formation plotted against valence numbers in FKBP and FRB constructs. *F*_N_ represents valence number of cytoYF_xN_, whereas *R*_M_ represents cytoCR_xM_. **(c)** Probability of iPOLYMER formation plotted against the total valance number N+M. In order to avoid bias, the combinations (N=1, M>1) and (N>1,M=1) were excluded from the data. Note that peptides with single valency should not lead to network formation, confirmed by the rare puncta formation in (b) for F1 or R1.

### Targeting puncta to subcellular compartments by signal sequences

By examining the expression patterns of YF_x1_ through YF_x5_, and CR_x1_ through CR_x5_, we noticed an increasing degree of nuclear localization as the valence number increased (Supplementary Fig. 3). This also led to rapamycin-induced puncta preferentially forming in the nucleus to cytosol, most likely due to the higher protein concentration in the nucleus. To direct the subcellular localization from the nucleus to the cytosol, we introduced a nuclear export sequence (NES) from mitogen-activated protein kinase kinase into the N-terminus of YF_xN_ and CR_xM_. These NES-tagged proteins, termed cytoYF_xN_ and cytoCR_xM_, were relatively abundant in the cytosol (Supplementary Fig. 3), and shifted the localization of rapamycin-triggered puncta towards the cytosol accordingly (Supplementary Fig. 4), though the effect was modest and the peptides were still prone to the nucleus with increased valency. A similar localization strategy was used to direct puncta formation to the plasma membrane by introducing a targeting sequence, such as a short plasma membrane-targeting peptide from a Lyn kinase into CR_x5_ (Lyn-CR_x5_). With this construct and mCherry-FKBP_x5_, rapamycin addition triggered puncta formation at the plasma membrane, which was verified by total internal reflection microscopy (Supplementary Fig. 5). In the following experiments, we mostly used the cytosolic versions, cytoYF_x5_ and cytoCR_x5_, but qualitatively similar results were obtained with the original YF_x5_ and CR_x5_.

### Valence number dependency of iPOLYMER puncta formation in computational model

Previous works ^28,36^, as well as the *in silico* experiments discussed above, clearly indicate that the valence number of binding molecules is a key factor for efficient aggregate formation. To explore the effects of valence numbers, we first carried out FRET measurement in cells co-transfected by the lower valence number constructs, YF_xN_ and CR_xM_ (N=M=1,2,3,4), and observed noticeable differences in the FRET kinetics upon rapamycin addition (Supplementary Fig. 1).

We further explored the effect of the valence numbers on the probability of puncta formation by testing all 25 different pairs (cytoYF_xN_,cytoCR_xM_), for *N, M* = 1,2,3,4,5. At small total valence numbers (i.e., when N+M ≤ 5), less than 15% of cells formed puncta (Fig. 3b and Fig. 3c). By contrast, the percentage of cells with puncta increased rather dramatically at large total valence numbers. The similarity of the dependency of aggregate formation on valence numbers in living cells (Fig. 3b) to that *in silico* (Fig. 2b) strongly suggests that the puncta formation *in situ* is indeed due to the FKBP/FRB polymer networks undergoing a phase transition. We could thus observe the dependency of both puncta formation kinetics and efficiency on valence number, which coincided with our expectation drawn from our theoretical considerations.

### Biophysical characterization of iPOLYMER puncta

To assess biophysical properties of the rapamycin-induced puncta generated in living cells, we performed fluorescent recovery after photobleach (FRAP) analysis of the components of rapamycin-induced puncta to examine their degree of remodeling. Cells expressing cytoYF_x5_ and cytoCR_x5_ were treated with rapamycin for 60 min and YFP was subsequently photobleached. The fluorescence recovery was significantly slower at the puncta as compared to a given cytosolic area with no apparent puncta (half-recovery time was 24 ± 2.5 s at the puncta and 11 ± 1.6 s outside the puncta) (Fig. 4a). Further fluorescence recovery was not observed even with an extended follow up, leaving a 77% immobile fraction (Fig. 4a and Supplementary Fig. 6a). The high immobile fraction suggests that cytoYF_x5_ is rather static inside the puncta, consistent with our previous observation that FKBP/FRB binding is practically irreversible ^37^. Interestingly, the rate of recovery within the cytosolic region outside the puncta was also slowed down after rapamycin treatment as compared to the case of cytosol without rapamycin (half-recovery time of 5.2 ± 0.84 s) (Fig. 4a), likely caused by small aggregates below the detection limit of our fluorescence microscope. To characterize the remodeling kinetics within the individual puncta, we performed a spot-bleaching of YF_x5_ within a single punctum 24 hours after rapamycin administration, which had developed to micrometer-size. As a result, we did not observe any striking fluorescence recovery in the bleached spot at least within one minute timeframe (Supplementary Fig. 6b).

**Figure 4.**
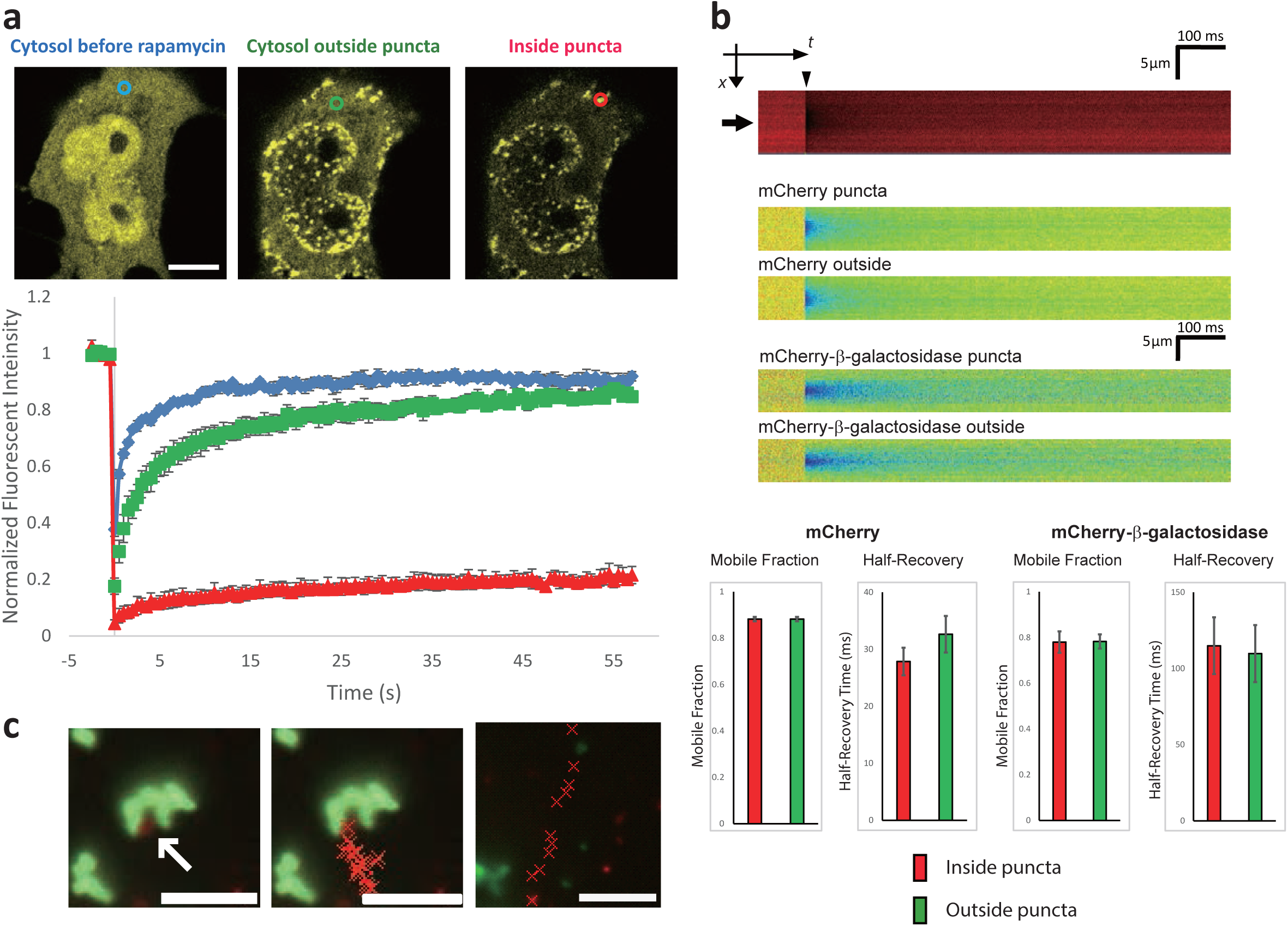
Biophysical analysis of iPOLYMER in living cells. **(a)** Top panel: Confocal fluorescence images of representative regions of cells expressing YF_x5_ and CR _x5_ subjected to FRAP analysis. Photobleaching was conducted under the following conditions: cytosolic region before administrating 333 nM rapamycin (cyan circle), cytosol outside the puncta (green circle), and inside the puncta (red circle). Scale bar: 10 μm. Lower panel: Fluorescent intensity transients before and after photobleaching. The colors correspond to those of the representative regions in the top panel. **(b)** iPOLYMER puncta allowed protein tracers to pass through. Top panel: Fluorescence intensity profile of mCherry in the cytosol in a line-scan FRAP experiment is shown in an *x-t* presentation. The fluorescence was photobleached at a single spot located in the middle of the puncta, as indicated by an arrow. The arrowhead indicates the time of bleaching. Middle panel: Representative normalized fluorescent intensity profiles shown for each experimental condition as a pseudo-colored image. Fluorescence recovery kinetics quantified from the data for mCherry and mCherry-β- galactosidase signals inside (puncta) or outside (outside) the puncta. Lower panel: The recovery kinetics were quantified by two parameters; mobile fraction (left graph) and half-recovery time (right graph). The values of these parameters were not significantly different inside and outside the puncta (p-value > 0.05, error bars: S.E.M.) for both tracer molecules. **(c)** Representative images of a mCherry-TGN38 labeled vesicle in contact with iPOLYMER punctum, indicated by the white arrow (left panel). The red crosses in the middle and right panels mark the position of the vesicle during a period of 150 s at 5 s intervals when colliding with the punctum (middle panel) and in the cytosolic region free of visible puncta (right panel). Scale bars: 5 μm.

### Molecules but not vesicles pass through iPOLYMER puncta

One of the essential features of hydrogels in terms of their biological function is their molecular sieving property ^30^, which enables them to play a role as a size-dependent filter. To test if the iPOLYMER induced puncta in living cells retain such property, we performed FRAP at the puncta, but this time against fluorescently-labeled diffusion probes, such as the red fluorescent protein mCherry and its fusion proteins of different sizes. First, mCherry alone was transfected in cells with cytoYF_x5_ and cytoCR_x5_. Following rapamycin addition, mCherry was photobleached at a single spot within the puncta (Supplementary Fig. 7a). The subsequent fluorescence recovery was quantified as normalized fluorescence intensity at the spot and the kinetics was analyzed (Materials and Methods, Supplementary Fig. 7b,c). To achieve sufficiently high spatio-temporal resolution, the FRAP experiment was carried out 24 h after rapamycin administration, when iPOLYMER puncta were typically grown to several micrometers in size (Supplementary Figs. 6b, 7a, and 8). The kinetics of subsequent fluorescence recovery of mCherry at the puncta were similar to that elesewhere (Fig. 4b, Supplementary 9a). We then repeated the experiment using a larger diffusion probe, namely mCherry fused to β-galactosidase that forms a tetramer complex with a diameter that is roughly 16 nm ^38^. The FRAP dynamics of mCherry-β-galactosidase were also similar to that observed outside the puncta (Fig. 4b, Supplemenatary Fig. 9b). Since the probe diameter is roughly 16 nm, which is greater than that of most soluble solitary proteins ^38^, we reasoned that proteins generally pass through the puncta without significant filtering effects. However, it was not clear if larger entities in cells, such as lipid vesicles, can pass through the puncta. To test this, we examined the movement of trans-Golgi vesicles, visualized with mCherry-TGN38 expression. Accordingly, we found that movement of mCherry-labeled vesicles was significantly slowed down upon hitting the puncta (24 ± 0.090 nm/s before hitting the puncta and 5.8 ± 0.011 nm/s after hitting the puncta, Fig. 4c), while there was no single event observed in which the vesicles penetrated through the puncta. Taken together, these intracellular diffusion assays implied that iPOLYMER-generated puncta in living cells retain a molecular sieving property, allowing most of soluble proteins to pass through, but disallowing penetration of larger entities, such as vesicles. Based on our observations, we estimated the effective pore size to be in the range of 16-70 nm, since the TGN38-labeled vesicles are reported to vary in size with a diameter around 70-140 nm ^39^.

### Ultrastructural analysis of iPOLYMER puncta and stress granules

The results described above imply the iPOLYMER puncta are functioning as a molecular sieve, supporting their character as a hydrogel. We therefore sought to compare the morphological nature of iPOLYMER puncta to that of a cytosolic protein-based structure with a hydrogel-like property. A stress granule is a protein-RNA assembly that is thought to undergo sol-gel-like phase transition upon various stress conditions ^40^ whose ultrastructure has been reported by electron microscopy (EM) ^41^. We used correlative electron microscopy (correlative EM) to compare the morphology of iPOLYMER puncta with that of stress granules (see Materials and Methods). Electron-dense, fibrillo-granular structure was observed at the iPOLYMER puncta (Fig. 5). These structures lack apparent membrane and resemble those of the actual stress granules morphologically (Fig. 5). These results suggest that the iPOLYMER puncta formed in living cell are likely similar to physiological hydrogel-like structures encountered *in situ.*

**Figure 5.**
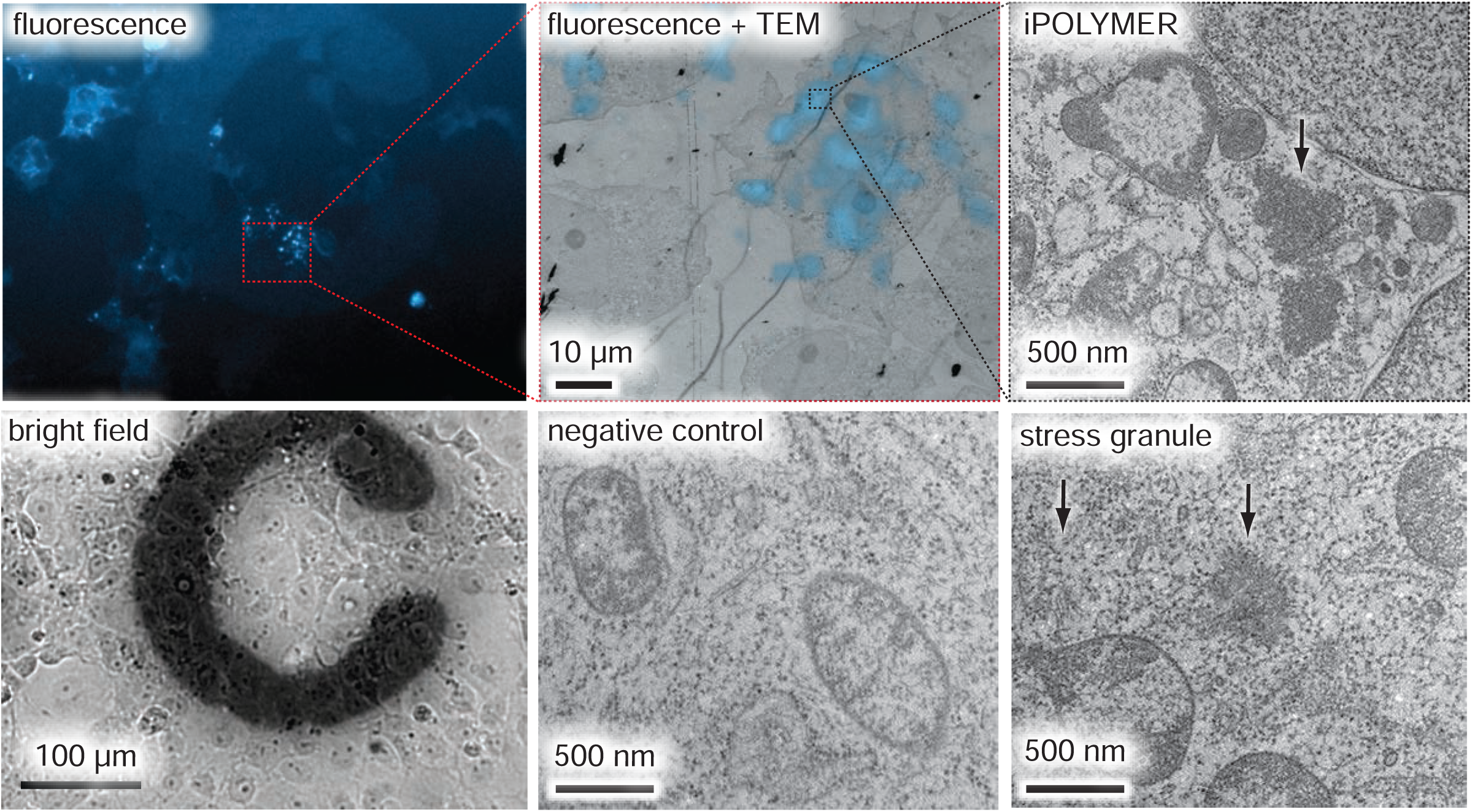
Correlative EM analysis of iPOLYMER puncta in living cells in comparison with stress granules. Transmitted EM (TEM) images of iPOLYMER puncta were obtained in COS-7 cells by correlating CFP-FRB_x5_ fluorescence image (top left panel (fluorescence)) with TEM image (top middle panel, shown as overlaid with correlated with fluorescence image (fluorescence + TEM), scale bar: 10 μm). The cells with apparent iPOLYMER puncta induced by 333nM rapamycin administraton were identified before EM imaging by referring to the grid pattern in the bright field image (bottom left panel (bright field), scale bar: 100 μm) High-magnification image of an iPOLYMER punctum is shown in the top right panel (iPOLYMER, scale bar: 500 nm), compared to the TEM image of an actual stress granule induced by 30 min incubation with 0.5 mM arsenite (bottom right panel (stress granule), scale bar: 500 nm). Negative control EM image of the cytosol without stress granule induction is also shown in the bottom middle panel (negative control). The iPOLYMER punctum exhibited an electron-dense granulo-fibrillar structure (black arrow in the top right panel) without any membranes surrounding it, resembling the actual stress granule (black arrows in the bottom right panel).

### *In vitro* reconstitution of iPOLYMER aggregate demonstrate hydrogel properties

To further explore the properties of hydrogel-like punctate material synthesized by iPOLYMER, we purified YF_x5_ and CR_x5_ with the aim to reconstitute rapamycin-induced hydrogels *in vitro.* When YF_x5_ and CR_x5_ were mixed at a final concentration that was lower than 5 μM, no puncta were observed under a fluorescence microscope. In contrast, when the protein concentration exceeded 5 μM, small puncta were observed after adding 1 μM of rapamycin (Supplementary Fig. 10). The puncta were enriched with fluorescence of both wavelengths and exhibited an elevated FRET efficiency. When the concentrations were increased to 100 μM for both peptides, and 500 μM for rapamycin, the solution became instantaneously turbid upon mixing (Fig. 6a), containing dense and irregularly shaped puncta when observed under the microscope (Fig. 6b). These puncta were rarely observed under negative controls in which DMSO was added instead of rapamycin, or rapamycin was added in the absence of CR_x5_ (Fig. 6b). Confocal imaging revealed the interior structure of the puncta to be irregular and heterogeneous (Supplementary Fig. 11). To demonstrate its structural integrity and its ability to hold water, we spun down the puncta by centrifuge, resulting in a clearly colored pellet (Fig. 6c). Inspection with a dissection microscope revealed that this pellet was structurally stable and optically translucent, with an elastic response to deformation (Fig. 6c, Supplementary Movie 7). Removal of the ambient solution did not result in an immediate loss of the pellet's shape or color (Fig. 6c), suggesting an ability of the material to absorb and retain water. Weighing the material before and after drying confirmed the water content calculated to be at least 75.2% (Supplementary Fig. 12). We therefore concluded that the rapamycin-induced aggregates of YF_x5_ and CR_x5_ were indeed hydrogels.

**Figure 6.**
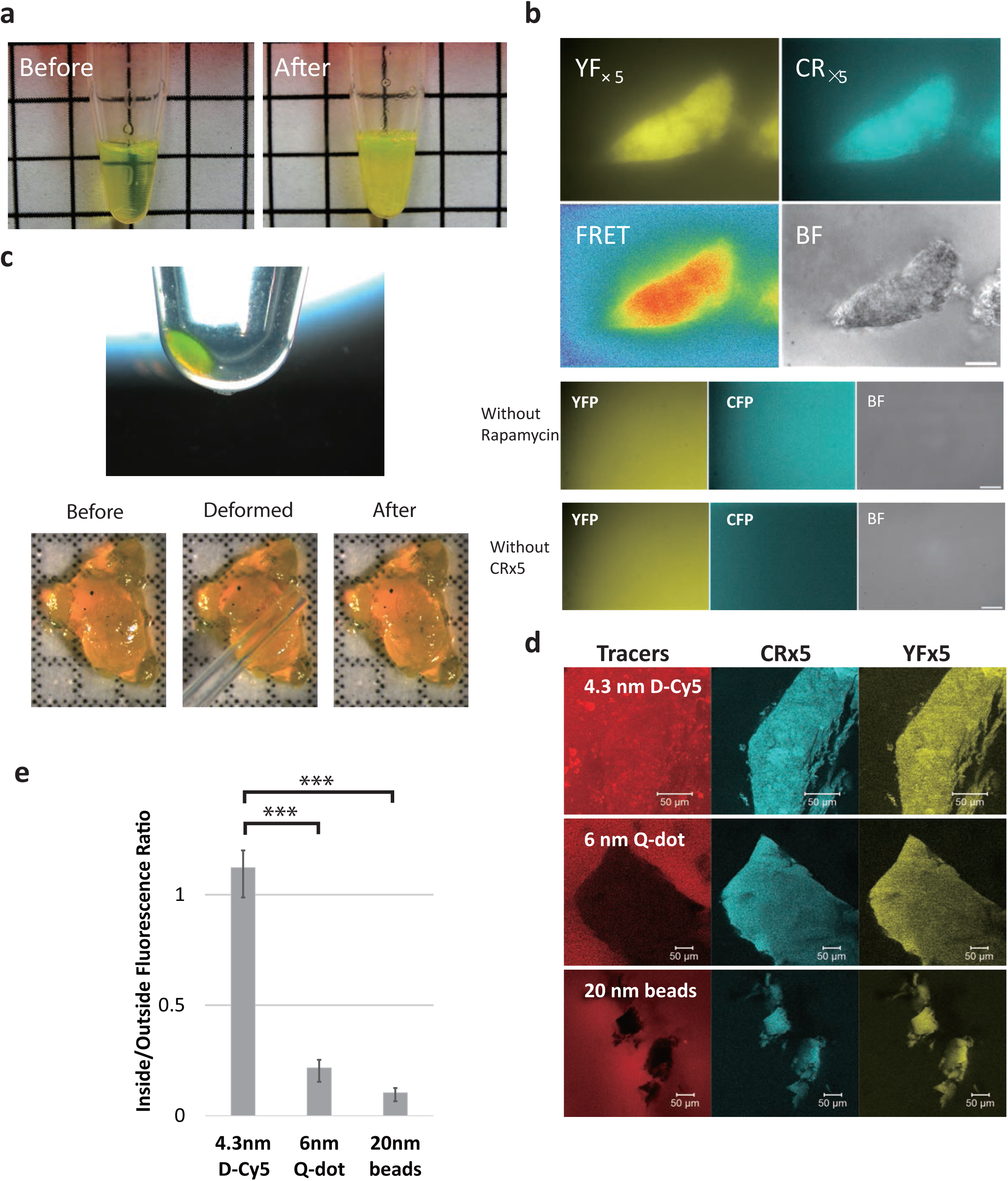
*In vitro* characterization of iPOLYMER. **(a)** Mixing 100 μM YF_x5_ and 100 μM CR_x5_ (left) with 500 μM rapamycin in a 1.5 mL tube instantly led to a turbid appearance (right). Size of grid: 0.5 cm. **(b)** Top panel: Fluorescent, FRET, and bright field (BF) microscopic images of iPOLYMER aggregates formed *in vitro.* Scale bar: 20 μm. Lower panel: Mixing 100 μM YF_x5_ with 100 μM CR_x5_ and DMSO did not form any aggregates (without rapamycin), and the same was true when mixing 100 μM YF_x5_ with 500 μM rapamycin (without CR_x5_). **(c)** Aggregates were collected by centrifuge and observed under a dissection microscope. Colored pellet was observed after centrifugation and removal of the supernatant (top panel). The pellet was isolated on a coverslip for further observation and experimentation (lower panel). The fragments were translucent with clearly defined shapes (before), and the aggregates retained their three-dimensional shape and translucent appearance after the removal, demonstrating the identity as a hydrogel. The hydrogel was mechanically deformed with a micropipette tip (deformed), and was confirmed to regain its original shape (after), which was almost indistinguishable from that before applying the deformation (before). **(d)** Pore size evaluation of the iPOLYMER hydrogel *in vitro.* Hydrogels collected by centrifuge were resuspended with fluorescent tracers with distinct diameters, and observed under a confocal fluorescent microscope. While a D-Cy5 tracer with 4.3 nm diameter penetrated into the hydrogel (top panel), 6nm-diameter Q-dot (middle panel) and 20 nm-diameter fluorescent beads (lower panel) were clearly excluded from the hydrogel. **(e)** The ratio of the tracer fluorescence intensity inside the hydrogel to that of outside the gel for each tracer molecule. Error bars, S.E.M.. ***, p<0.01. The observed difference of the ratios associated with D-Cy5 (4.3 nm) and Q-dot (6 nm) suggest that the hydrogel functions as a molecular sieve with an effective pore size of 4.3-6 nm.

### iPOLYMER hydrogel as a size-dependent molecular sieve

To assess the molecular sieving potential of the hydrogel *in vitro,* we performed diffusion assays using fluorescent diffusion tracers of varying sizes. We infused these tracers one at a time into a solution containing rapamycin-induced hydrogel (using a spun down mixture of 100 μM YF_x5_, 100 μM CR_x5_, and 500 μM rapamycin), and measured the ratio of fluorescence intensity inside and outside the hydrogel under a confocal microscope (Fig. 6d). More specifically, we employed the following diffusion tracers that are known to exhibit minimal adsorption to general proteins: D-Cy5 (4.3 nm in diameter) ^42^, CdSeS/ZnS alloyed quantum dot (Q-dot) (6 nm in diameter), and fluorescent beads (20 nm in diameter). While the 4.3-nm tracer did penetrate into the hydrogel almost freely, the 6-nm and 20-nm tracers were clearly excluded from the hydrogel (Fig. 6d, Supplementary Fig. 13). We thus estimated the effective pore size of the hydrogel *in vitro* as 4.3-6 nm.

### Functionalizing iPOLYMER gels

Since intracellular hydrogel-like biomaterial has been proposed to constitute a functional platform for cellular activities, we next aimed to utilize iPOLYMER hydrogels as a scaffold to rapidly nucleate a biological entity. We expected that the iPOLYMER paradigm can be effectively used to study RNA granules, due to their gel-like property along with their morphological resemblance to iPOLYMER puncta *in situ* (Fig. 5). As mentioned earlier, a stress granule is a protein-RNA assembly that undergoes sol-gel-like phase transition upon stress stimulus ^40^. Previous studies have shown that one stress granule component, TIA-1, self-assembles upon treatment with arsenate as a stress, turning into a stress granule ^43,44^. TIA-1 consists of two domains: an RNA recognition motif (RRM) that binds to polyadenine (Poly-A) containing RNAs, and a prion-related domain (PRD), which self-assembles into a gel-like state. It was found that when the RRM domain is fused with an exogenous aggregation- promoting domain in place of the original PRD, RNA granules would form spontaneously ^43^. To reproduce the formation of RNA granules in an inducible manner, we first replaced the PRD domain with one of the iPOLYMER components, CR_x5_, to produce a fusion protein RRM-CR_x5_. We then induced the formation of intracellular hydrogels with coexpressed cytoYF_x5_ (Fig. 7a). After administering rapamycin for 1h, cells were immunostained for endogeneous PABP-1, an RNA-binding protein known to accumulate in stress granules. The iPOLYMER puncta formed with RRM-CR_x5_ were colocalized with endogenous PABP-1 (Fig. 7b). This suggested that the functionalized iPOLYMER puncta, or the stress granule analogues, could sequester poly-A containing mRNAs in a similar fashion as the actual stress graunles. In contrast, PABP-1 was not accumulated in iPOLYMER puncta without functionalization with TIA-1 RRM (Figure 7b).

**Figure 7.**
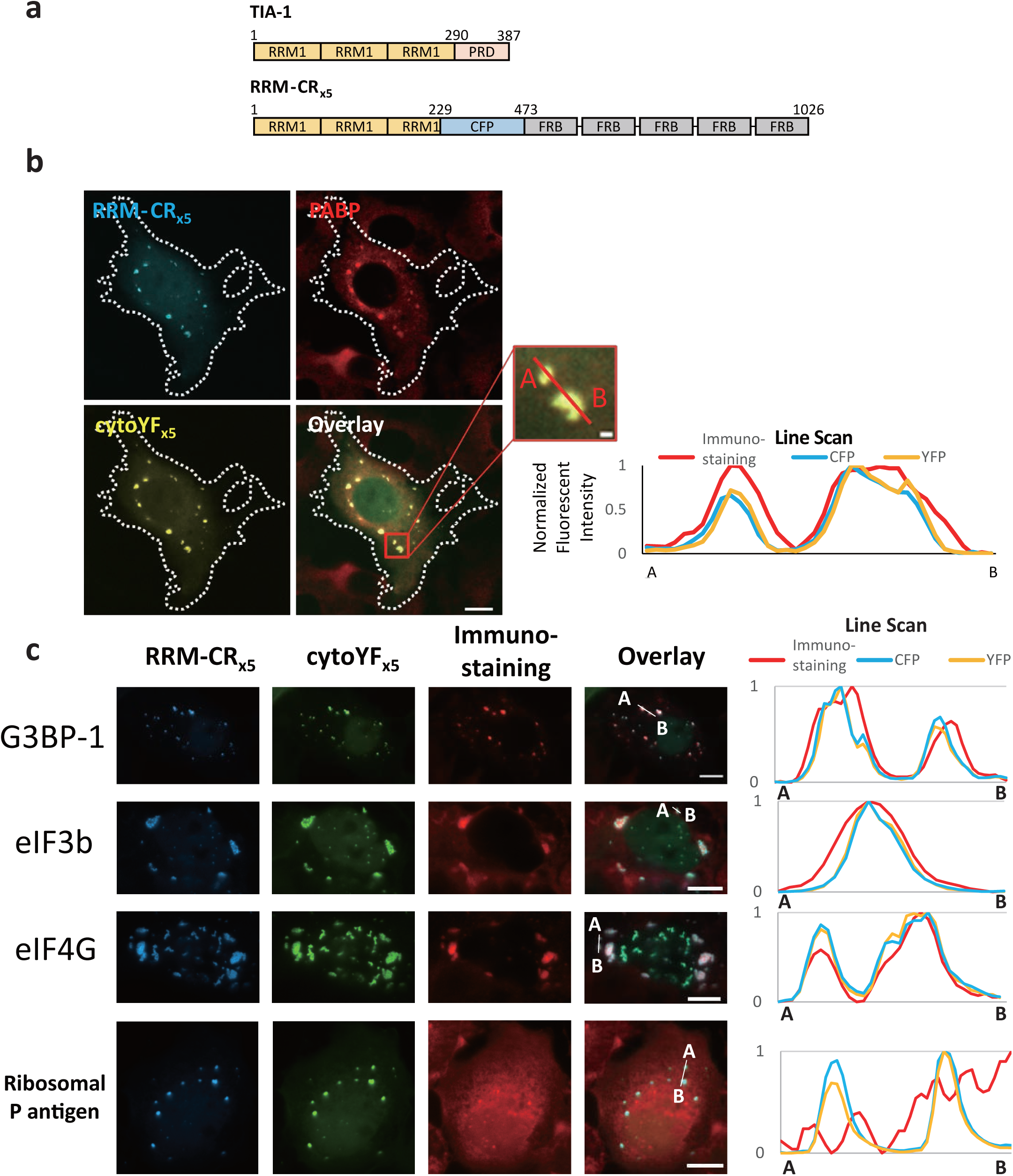
Reconstituting RNA granules by using iPOLYMER as scaffold. **(a)** Schematic illustration of the RRM-CR_x5_ construct used in RNA granule reconstitution. Three RRM domains from TIA-1 were fused to CR_x5_. **(b)** Immuno-staining images of COS-7 cells expressing RRM-CR_x5_ and cytoYF_x5_ treated with rapamycin to form iPOLYMER puncta. Scale bar: 10 μm. The line scan plot from A to B in the enlarged image confirms co-localization of endogenous PABP with the functionalized iPOLYMER puncta. **(c)** Immuno-staining of the RRM-functionalized iPOLYMER with the universal stress granule markers G3BP-1, eIF3b, and eIF4G and corresponding line scan plots from A to B shown in each overlay image. As a negative control, ribosomal P antigen, which does not accumulate in the stress granules, was also stained. RRM-functionalized synthetic analogue of RNA granules accumulated all the three stress granule markers (7/17 cells showed accumulation of G3BP-1, 12/29 cells of eIF3b, and 8/15 cells of eIF4G), while ribosomal P antigen accumulation was not observed (0/27 cells).

### Is gelation required for recruitment of mRNAs?

We evaluated if the recruitment of mRNAs to the stress granule analogues requires gelation. To test this, we tried to concentrate RRM at the lysosomal membranes without gelation, intending to achieve both RRM density and the size similar to those of the cytosolic iPOLYMER gels. We added rapamycin and recruited RRM-CR_x5_ to LAMP-YF that was expressed at the cytosolic face of lysosomes ^45^. We chose lysosomes as they have similar microscopic length scale (i.e., < a micron). Such a condition should not experience sol-gel transition due to a lack of multivalent associations, and this was indeed the case; neither exogenous nor endogenous PABP-1 co-localized with lysosomes (Supplementary Fig. 14). We then addressed if the failed PABP-1 accumulation was due to a lack of the sol-gel transition and/or to a local environment specific to the lysosomal membranes compared to the cytosol. For this purpose, we repeated the experiment now with LAMP-YF_x5_, which contains five tandem FKBPs co-expressed with RRM-CR_x5_, and we did not observe PABP-1 accumulation or puncta formation at the lysosomes (Supplementary Fig. 14). These results suggest that PABP-1 accumulation at the RRM-functionalized iPOLYMER puncta may require the cytosolic environment that is distinct from that on the cytosolic surface of lysosomal membrane. It should be noted, however, that we do not have a feasible method to directly detect the existence of hydrogel-like phase in the experiment, thus cannot exclude the possibility that hydrogelation plays a role in PABP-1 accumulation.

### Characterization of the iPOLYMER-based stress granule analogues

To further characterize the iPOLYMER-based synthetic stress granule analogues (i.e., the RRM(TIA-1)-functionalized iPOLYMER) in more details, we performed immunostaining against three universal markers of physiological stress granules: G3BP1, eIF4G, and eIF4b. These are mRNA-associated proteins that are present in stress granules induced by various stresses. In addition to these positive markers, we looked at a ribosomal large subunit P antigen, a protein that also binds to mRNAs and regulates translation. Unlike the previous three RNA binding proteins, the ribosomal P antigen does not accumulate in stress granules, and it is this absence that is thought to be responsible for a critical function of stress granules, i.e., a halted translation of homeostatic proteins ^46^,^47^ (Supplementary Fig. 15). We generated the stress granule analogues using RRM-CR_x5_ and cytoYF_x5_, as described earlier. A subsequent immunostaining against the four proteins revealed colocalization of the three stress granule markers, but not of ribosomal P antigen, with the stress graunle analogues (Fig. 7c). The accumulation of G3BP1, eIF4G, and eIF4b at the analogues was dependent on the presence of both rapamycin and RRM, as it was abolished when repeating the experiment either without rapamycin administration, or using cytoCR_x5_ that lacks RRM instead of RRM-CR_x5_ (Supplementary Fig. 16). These results demonstrated that the stress granule analogues generated by iPOLYMER retain a biochemical profile similar to that of physiological stress granules.

### iPOLYMER principle to achieve reversible protein aggregates; development of iPOLYMER-LI

Although iPOLYMER-based analogues of stress granules reproduced the selectivity of their molecular components, the analogues still differ from their physiological counterpart in several key properties. Irreversibility in granule formation, which is due to the irreversible nature of rapamycin-based CID, would be one of the most striking differences. Also, the reversible and dynamic nature of stress granule is considered to be an essential feature of the organelle, possibly contributing to its biological function^xx^. The irreversible remodeling within the rapamycin-based iPOLYMER puncta could limit the application of the technique in broader spectrum of biological events. To advance the general applicability of iPOLYMER, we applied the same design principle, i.e., multivalent dimerizer peptides, to light-inducible dimerizers. Unlike rapamycin-based CID, light-inducible dimerizers are often reversible. More specifically, we used iLID and SSPB, which dimerize upon irradiation with blue light ^48^. We synthesized six tandem iLIDs and six tandem SSPBs, which were fused to YFP and mCherry, respectively (Fig. 8a). When co-expressed in cells, these two multivalent peptides formed cytosolic aggregates upon light illumination at 488 nm (Fig. 8b). The aggregates gradually disappeared over ten minutes after stopping the blue light illumination, representing the reversible induction of protein aggregates (Fig. 8b). The reversible aggregates could also be induced subcellularly and repetitively (Fig. 8c, Supplementary Video 7). By adapting optogenetics, we could demonstrate reversible iPOLYMER, iPOLYMER-LI, for <u>i</u>ntracellular <u>p</u>roduction <u>o</u>f <u>l</u>ight-<u>y</u>ielded <u>m</u>ultivalent <u>e</u>nhancers with <u>l</u>ight-<u>i</u>nducibility, armed with improved temporal and spatial control.

**Figure 8.**
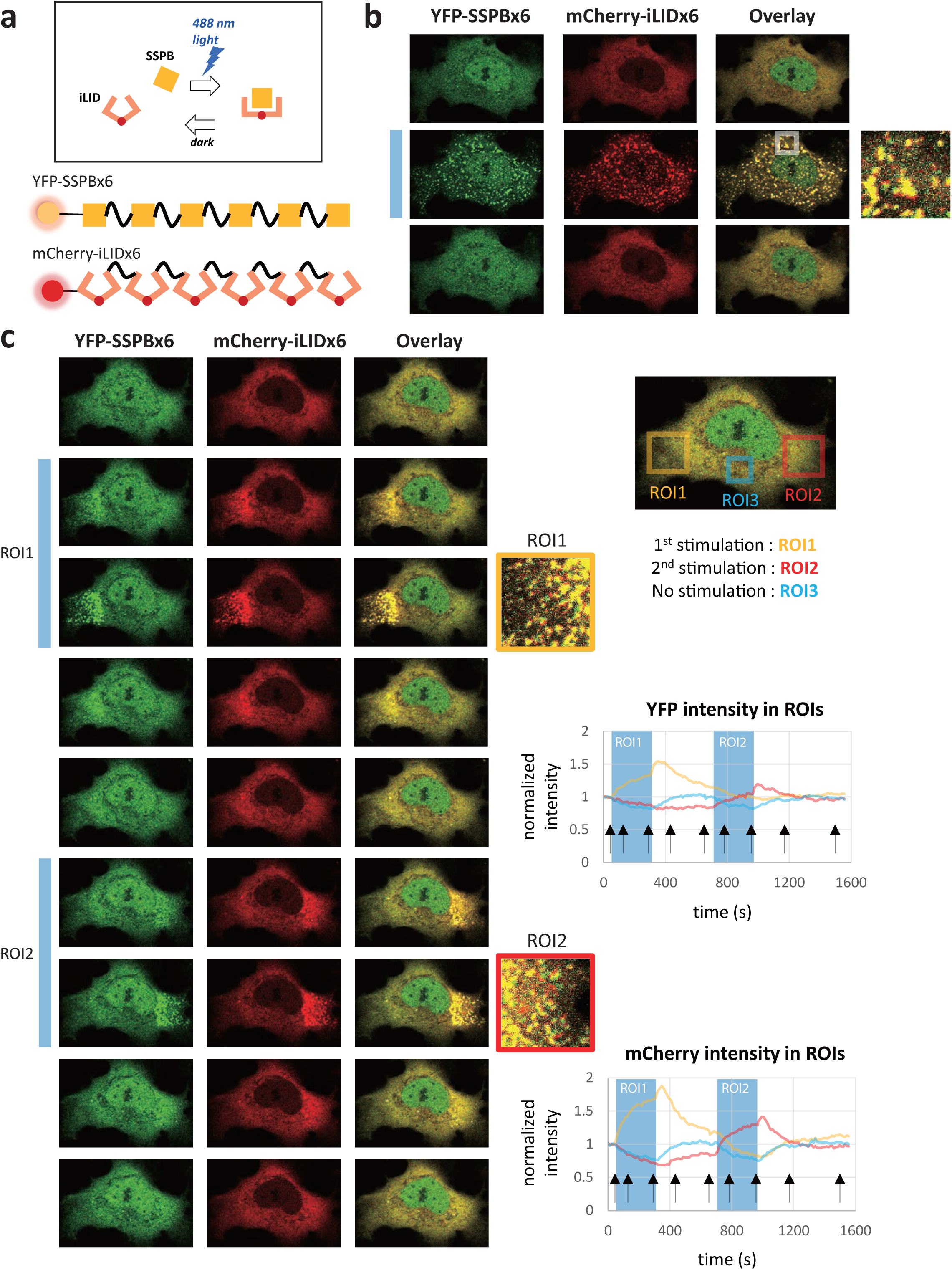
Light-inducible iPOLYMER-LI provides reversibility and spatial control over puncta formation. **(a)** Designs of light-inducible iPOLYMER-LI constructs. The two peptides, SSPB and iLID, bind to each other upon blue light (488 nm) stimulation in a reversible manner (top panel). YFP-SSPB_x6_ and mCherry-iLID_x6_ contain six repeats of iLID and SSPB, respectively, spaced by nine amino acid linker sequences (lower panel). Due to design principles similar to YF_x5_ and CR_x5_, the two peptide chains are expected to form a polymer network upon light irradiation. However, unlike YF_x5_ and CR_x5_, the network formation is reversible. **(b)** Reversible puncta formation by YFP-SSPB_x6_ and mCherry-iLID_x6_. The cell was irradiated with 488 nm laser before each frame during the stimulus. The blue rectangle labels images observed under stimulus. Light-induced formation of the protein aggregates was readily observed within 15 min (middle panel and magnified detail). By ceasing stimulation, the aggregates were dispersed within 7.5 min (lower panel), demonstrating the reversible nature of the light-inducible version of iPOLYMER, iPOLYMER-LI. **(c)** Aggregates were formed repetitively at distinct locations within the same cell using the iPOLYMER-LI (left panel). The blue rectangles label images obtained under stimulation. Magnified views of two stimulated regions of interest, ROI1 and ROI2, are also shown highlighted by colors that correspond to those in the right top panel. Fluorescence intensities are shown in the right middle and right lower panels for YFP and mCherry respectively, in which the stimulation timing is highlighted by blue. During the first stimulation, only ROI1 was illuminated, whereas only ROI2 was illuminated during the second stimulation. The apparent overshoot in fluorescence intensity right after each stimulation is probably an artifact, since the fluorescence intensities at the puncta were often saturated during the stimulation. Black arrows in the plots indicate the timings of the images shown in the left. Taken together, these results show that fluorescent puncta were dynamically formed and dispersed locally, demonstrating both the reversibility and spatio-temporal control over this process.

**Figure 9.**
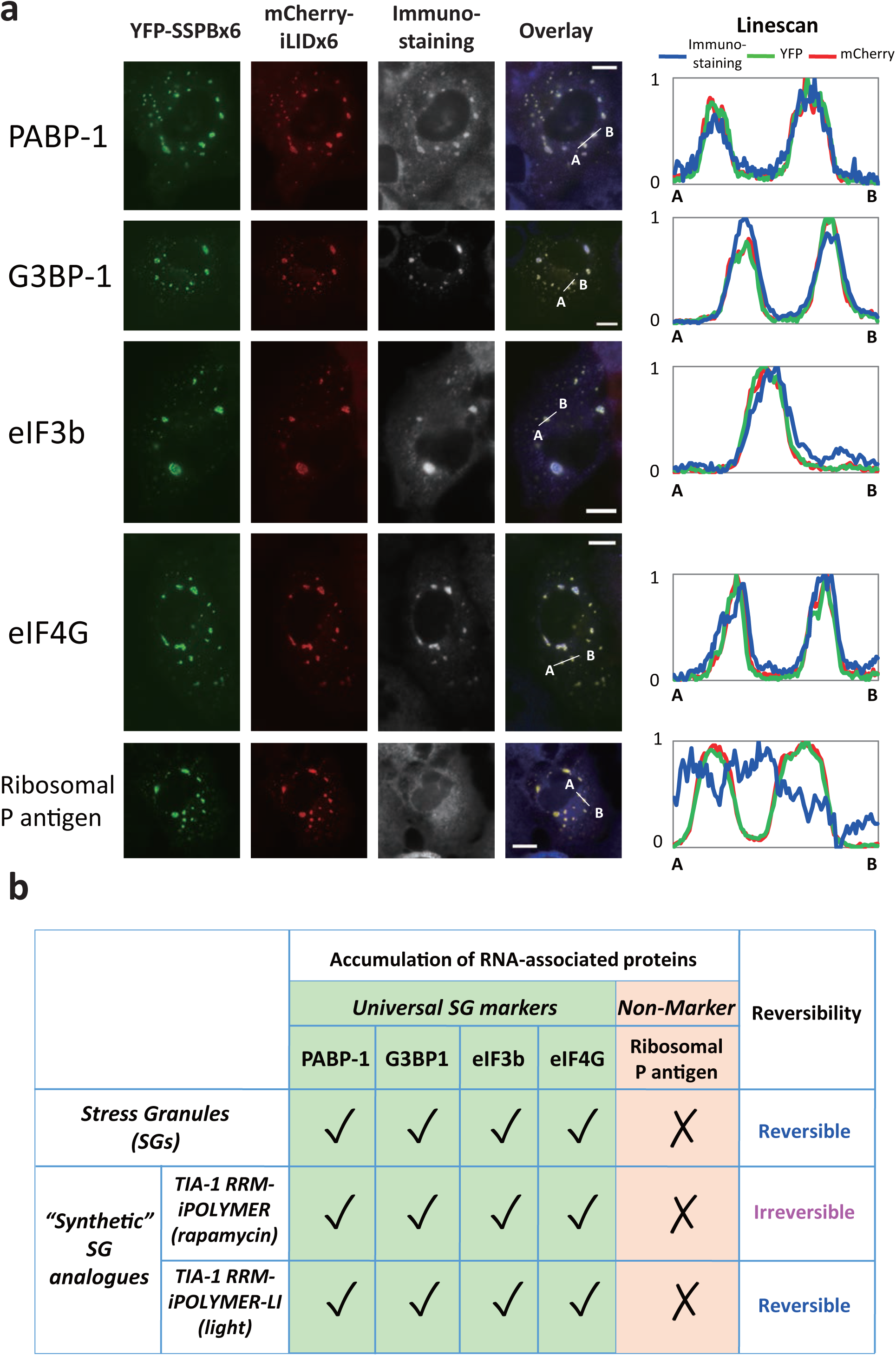
RNA granule reconstitution by light-inducible iPOLYMER-LI. **(a)** Immuno-staining of the RRM-functionalized iPOLYMER-LI with the universal stress granule markers PABP-1, G3BP-1, eIF3b, and eIF4G and corresponding line scan plots from A to B shown in each overlay image. Immunostaining results are shown in blue in overlay images. As a negative control, ribosomal P antigen, which does not accumulate in the stress granules, was also stained. RRM-functionalized iPOLYMER-LI puncta (YFP-SSPB_x6_ and mCherry-iLID_x6_) accumulated all the three stress granule markers (33/65 cells showed accumulation of PABP-1, 25/40 cells of G3BP-1, 23/51 cells of eIF3b, and 52/75 cells of eIF4G), while ribosomal P antigen accumulation was not observed (0/75 cells). Scale bars: 10 μm. **(b)** Comparison between iPOLYMER-and iPOLYMER-LI-based stress granule analogues and the actual stress granules. Stress granule analogues were similar to its physiological counterparts in the specific recruitment of universal stress granule markers. Rapamycin-induced analogues differ from physiological stress granules in reversibility of the formation process, while iPOLYMER-LI-based analogues have overcome the discrepancy.

### Reversible iPOLYMER-LI successfully produced stress granule analogues

The novel class of iPOLYMER, iPOLYMER-LI, can produce protein-based aggregates in living cells in a reversible manner. However, the ability of iPOLYMER-LI in reconstituting stress granules in cells has yet to be demonstrated. We therefore fused RRM domains from TIA-1 to the N-termini of the two iPOLYMER-LI peptide chains to make functionalized peptides, TIA1 RRM-mCherry-iLIDx6 and TIA1 RRM-YFP-SSPBx6. We first produced iPOLYMER-LI puncta with TIA1 RRM-mCherry-iLIDx6 and YFP-SSPBx6 by illuminating cells with blue light for 1 hr, performed immunostaining against stress granule markers, and found little significant accumulation of any of the markers at the puncta. We suspected that the relatively low accumulation of RRM domains was the reason for the lack of accumulation. Therefore, we produced puncta by TIA1 RRM-mCherry-iLIDx6 and TIA1 RRM-YFP-SSPBx6, with RRM domains on both peptides. As a result, we confirmed accumulation of all four stress granule markers, PABP-1, G3BP-1, eIF3b, and eIF4G at the functionalized iPOLYMER-LI puncta, while we did not observe any accumulation of ribosomal P antigen. The accumulation was light stimulus-dependent and RRM domain-dependent (Supplementary Fig. 17). The selective colocalization of universal stress granule markers with functionalized iPOLYMER-LI puncta revealed that the reversible version of iPOLYMER produced stress granule analogues not only in terms of selective molecular components, but also of reversible mechanisms of puncta formation, which is one of the key properties of the organelle.

### Discussion

To overcome difficulties in enabling gel formation *in situ,* we first developed a realistic computational model of iPOLYMER to assess the effects of various parameters on gel formation and its properties. This presented us with a deeper understanding of the problem of gel synthesis, which rationally guided our experimental design and provided further validation of our experimental findings and conclusions. More specifically, our computational model exhibited valency-dependent polymer network formation with striking resemblance to our experimental results, whereas *in vitro* reconstitution of the polymer network confirmed the identity of the material as a hydrogel (for detailed comparison, see Discussion in Supplementary Methods). We also experimentally estimated the effective pore size of the hydrogels *in vitro* to be 4.3-6 nm. These sizes, however, were significantly smaller than the ones estimated in living cells (16-70 nm), a discrepancy that could be explained by differences in protein and rapamycin concentrations used in our experiments (proteins: 100 μM in solution, while typically less than 1 μM for overexpressed proteins in mammalian cells with a CMV promoter, rapamycin: 1-500 μM *in vitro,* while typically 333 nM in extracellular solution for cells). It is very likely that these differences result in hydrogels with different network densities and thus with different pore sizes. In addition, an experimental procedure required concentration of the hydrogels by centrifugation during our *in vitro* evaluation, which may have affected the pore size as well.

In the current study, we took advantage of the iPOLYMER paradigm to induce formation of stress granule analogues in living cells. Although there have been several implications of stress granules as a hydrogel, it has recently been claimed that many non-membrane-bound organelles including RNA granules such as stress granules have a common dynamic feature often referred to as liquid droplet like ^9,13^. One report indeed indicates that a given low complexity domain can form hydrogels *in vitro,* while it shows liquid droplet-like behavior *in situ,* and that the molecular conformation is similar in both cases ^49^. However, how these distinct physical properties (static hydrogels vs. dynamic liquid droplets) emerge remains to be understood, and the iPOLYMER strategy may contribute to understanding of their structure and function in the future. Of note, design principles of iPOLYMER could dictate its functionality. Rapamycin-based iPOLYMER granules, for instance, exhibited the constant growth in size due to its irreversible nature, which may allow these granules to go beyond optimal length scale to be functional as an RNA granule. If this happens to be the case, optogenetic iPOLYMER-LI may be favored due to its reversible and precise operation in space and time. Also, we cannot fully exclude the possibility that iPOLYMER/iPOLYMER-LI puncta formation itself becomes a stress to cells and activate stress pathways, which may secondarily affect molecular features of the iPOLYMER-induced RNA granules.

Another promising application of the iPOLYMER would be reconstituting a nuclear pore complex (NPC), which utilizes a FG-repeat-based, reversible hydrogel for its selective permeability of molecules ^30^. Synthetically reconstituting the NPC-like permeability barrier based on the iPOLMER may result in an interesting insight into the NPC biology.

From a technical point of view, the iPOLYMER has several advantages over previous counterparts that can form protein-based aggregates in living cells ^50–54^. First, we are the first to determine the detailed physical nature of the aggregates, and concluded them to be a hydrogel. Second, puncta formation of iPOLYMER does not rely on protein unfolding or domains from pathogenic proteins, both of which possibly can trigger intrinsic cellular responses, unlike several recently reported techniques ^52,54^. Third, iPOLYMER provides a flexible, generalizable framework, as indicated by development of the optogenetic iPOLYMER-LI. Taken together, the iPOLYMER armed with spatio-temporal control could potentially become a powerful means to study granular structures in cells, and gel-like aggregates that are proposed to exhibit neurodegenerative toxicity ^55^.

## Materials and Methods

### Reagents

Rapamycin was purchased from LCLab and prepared as dimethylsulfoxide (DMSO) stock solutions. Fluorescently labeled ethylenediamine-core generation-4 hydroxyl-terminated poly(amidoamine) dendrimer, D-Cy5 ^42^, was kindly provided by Rangaramurajan M. Kannan's group. CdSeS/ZnS alloyed quantum dots with 665 nm emission wavelength were purchased from Sigma Aldrich. FluoSpheres Fluorescent Microspheres (fluorescent polystyrene beads) were purchased from Molecular Probes. Antibodies against G3BP, eIF3b, eIF4G, and ribosomal P antigen, and sodium arsenite were kind gifts from Nancy Kedersha.

### Computational modeling

*In silico* implementation of iPOLYMER was based on a realistic kinetic Monte Carlo simulation algorithm, which produced sufficiently accurate approximations of stochastic reaction-diffusion dynamics. Model details and computational analysis can be found in the Supplementary Methods.

The model employs a stochastic biochemical reaction system, which contains three types of molecules, FKBP, FRB and rapamycin. These molecules interact according to four reversible reactions (Fig. 2a) and are subject to random diffusion. The binding sites in FKBP and FRB were labeled as free or bound at each time point, along with the information of binding partners. At a given time, the system may contain a mixture of FKBP, FRB, and rapamycin molecules, as well as aggregate molecules formed by the mutual binding of these three basic molecules with other larger compound molecules.

To model iPOLYMER, we spatially discretized the well-known continuous-space Doi model of stochastic reaction-diffusion ^56,57^, and obtained a physically valid approximation based on the reaction-diffusion master equation (RDME) ^58–62^. This led to a Markov process model that describes the time evolution of the location of each basic or aggregate molecule at a resolution of one voxel in the system. We simulated the resulting process by a stochastic kinetic Monte Carlo algorithm. For our computational analysis, we modeled the CID system in a predefined volume of a subcellular size, discretized the system in each spatial direction resulting in a given number of equally sized voxels that satisfy the modeling assumptions and constraints, used experimentally verified kinetic rate values for certain reactions, and plausible values for the kinetic rates of the remaining reactions (see Supplementary Methods).

### DNA constructs

Construction of CR_xM_ and YF_xN_: To generate the YFP-FKBP (YF_X1_) and CFP-FRB (CR_x1_) constructs, two distinct polymerase chain reaction (PCR) products encoding FKBP and FRB were digested with BsrGI and XhoI and inserted into the pEYFP(C1) and pECFP(C1) constructs (Clontech), respectively. To prepare the YF_x2_ and CR_x2_ constructs, FKBP and FRB fragments with a 12 amino acid linker sequence (SAGGx3) were generated by PCR and cloned into YF_x1_ and CR_x1_ constructs, respectively. The same strategy was used to generate two series of multivalent FKBP and FRB constructs.

Construction of cytoCR_xM_ and cytoYF_xN_: Nuclear export signal from MAPKK was inserted into the Nhel and Agel cloning sites of CR_xM_ and YF_xN_, respectively.

Construction of YF_x5_-His and CR_x5_-His for purification: Coding sequences of CR_x5_ and YF_x5_ were amplified by PCR from the plasmids described above, respectively. The obtained fragments were then digested with NcoI and NotI cloned into the corresponding sites in pET28a.

Construction of RRM-CR_x5_ and RRM-CR_x5_ (pMT2): The YFP-TIA-1 and pMT2 vectors were kindly provided by Paul Anderson's group. The RRM domain was amplified from YFP-TIA-1 using 5' TAGCTAGCGCCACCATGGAGGACGAGATGCCC and 5’ATACCGGT-CCTAGTTGTTCTGTTAGCCCAGAAG. The PCR products were digested with NheI and AgeI, and cloned into the corresponding sites in CR_x5_ to obtain RRM-CR_x5_. RRM-CR_x5_ was then amplified by 5' ACGCCTGCAGGGCCACCATGGAGGACGAG 3’ and 5' ACGCAATTGTCAGTTATCTAGATC-CGGTGG 3’, and digested by SbfI and MfeI. The digested fragments were then cloned into the PstI and EcoRI cloning sites of the pMT2 vector to obtain TIA-1RRM-CR_x5_ (pMT2). Overexpression of many stress granule-associated proteins induces spontaneous stress granules in the absence of additional stress. It is known that pMT2 vector may express inhibitors of the eIF2α kinase PKR, which is required for the spontaneous stress granule formation ^46^. We therefore adopted pMT2 vector to inhibit excessive formation of stress granules in some experiments.

To make the light-inducible iPOLYMER constructs mCherry-iLID_x6_ and YFP-SSPB_x6_, we synthesized DNA sequence encoding three tandem iLID and SSPB (Genscript), and subcloned two tandem sequences of them in pmCherry(C1) and pEYFP(C1) vectors, respectively, between BglII and BamHI sites in the original vectors, taking advantage of the compatible ends cleaved by the two restriction enzymes.

### Cell culture and transfection

COS-7 cells were cultured in a DMEM (GIBCO) medium supplemented with 10% FBS and 1% Penicillin Streptomycin (Life Technologies) in a 37°C and 5% CO_2_ incubator. Cells were transiently transfected using either FugeneHD (Roche) or Amaxa Nucleofector (Lonza) and seeded in 8-well chamber slides (Thermo).

### Cellular imaging of iPOLYMER puncta formation

Cells were transfected with desired constructs, seeded in an 8-well chamber slides and incubated for 36-48 h. The culture medium was then washed and replaced with Dulbecco's Phosphate-Buffered Saline (Gibco) for imaging. Imaging was conducted with an Olympus inverted microscope and a Zeiss LSM 780 confocal imaging system. FRAP experiments were conducted on Zeiss LSM 780. All images were processed using Metamorph imaging software (Molecular Devices). For inducing hydrogel formaition by iPOLYMER, rapamycin in DMSO was diluted with DPBS to 3.33 μM, and added to the chamber by 1:10 dilution.

### Quantification of iPOLYMER puncta formation

COS-7 cells were simultaneously transfected with cytoCR_xM_ and cytoYF_xN_ and grown 36-48 h as described above. Rapamycin was then added (333 nM) and cells incubated 20 min at 37 °C with 5% CO_2_. Cells were then fixed with 4% (w/v) paraformaldehyde and imaged on a Zeiss Axiovert 135 TV microscope with a Qlclick camera (Qlmaging). For each cell imaged, a Z-stack of 5 planes separated by 1 μm was taken to ensure that any puncta present in the cell would be observed. Cells were considered to be punctated if at least one punctum with both CFP and YFP fluorescence was present, and to be non-punctuated if no such structure was present. The results are the mean ± S.E.M from three independent experiments.

### FRAP experiment and analysis

All FRAP experiments were carried out with Zeiss LSM780 confocal microscope using bleaching function. In the FRAP evaluation of the turnover rates of the iPOLYMER peptides, data was acquired by raster scanning. YFP was photobleached by 514 nm laser within a circular ROI for bleaching. The fluorescence intensity within the ROI was monitored, and was normalized to the intensity before bleaching. The normalized fluorescence recovery over time, *F*(t), was then fitted to an exponential function;

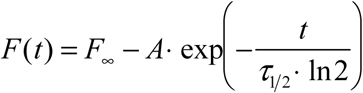
to obtain mobile fraction *F*_∞_, half-recovery time *τ*_1/2_, and *A* as fitted parameter values.

To monitor the diffusion of the tracer, mCherry or mCherry-β-galactosidase, within the iPOLYMER puncta, higher resolution was required in terms of both space and time. We therefore adopted line-scan data acquisition for higher frame rate, and spot-bleach method at a single spot without any scanning for photobleaching. Fluorescence recovery at the bleaching spot was monitored and analyzed by fitting the normalized fluorescence intensity transient *F*(*t*) to the following function.

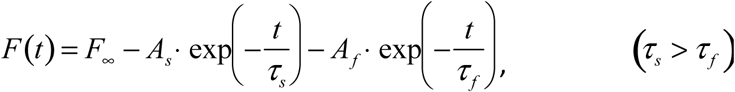
to obtain five fitting parameter values, *F_∞_*, *A_s_*, *A_f_*,*τ_s_*, and *τ_f_ F_∞_* is immobile fraction, *A_s_* and *A_f_* are amplitudes of slower and faster recovery component, respectively, and *τ_s_* and *τ_f_* are characteristic decay times for slower and faster component, respectively. In order to characterize the diffusion kinetics by a single parameter, we then computationally quantified the half recovery time, *τ*_1/2_ by interpolation (Supplementary Fig. 8c). Briefly, the fitted function was plotted with time step Δt = 0.1 ms. The time required for the half recovery was then calculated by linear interpolation between the plotted data points to obtain *τ*_1/2_. All the fittings were done on Igor Pro software (WaveMetrics), using built-in curve fitting formulas and functions.

### Protein purification and *in vitro* hydrogel formation

CR_x5_-His and YF_x5_-His vectors were transformed into the BL21(DE3) *Escherichia coli* host strain (Novagen). Protein expression was induced by 0.1 mM isopropyl β-D-1-thiogalactopyranoside and the proteins in the supernatant were bound to Ni-nitrilotriacetate resin (Qiagen). After elution by 250 mM imidazole, the eluted fraction was dialyzed and purified further by Mono Q 5/50 GL anion exchange column. Purified YF_x5_ and CR_x5_ were premixed in 20mM Tris pH8.5, 300mM NaCl. Subsequently, rapamycin stock in DMSO was added with a 1:100 dilution and immediately mixed vigorously. For the experiments associated with pore-size estimation, the solution was further centrifuged at 16,000 g for 30-60 min at room temperature. Pellets were suspended again in the solution by brief pipetting, they were transferred onto the coverslip, and the solution containing both fluorescent tracer and rapamycin was added on top. The penetration of the fluorescent tracers was examined by LSM780 confocal microscope (Zeiss).

### TGN38 vesicle movement analysis

Cells were transiently transfected with TGN38-mCherry with cytoYF_x5_ and cytoCR_x5_. Cells were first induced to form iPOLYMER for 1 h, then imaged while maintained at 37^o^C and 5% CO_2_. The images were taken every 5 sec. The velocity of the TGN38-posibive vesicles was estimated by dividing the distance that the vesicles traveled within the xy plane between two consecutive frames by the 5 sec interval. The images were analyzed using the “track points” function in a Metamorph image analysis software (Molecular Devices).

### Induction of physiological stress granules and their analogues

To induce stress granule formation, cells were incubated in 0.5 mM sodium arsenite-containing culture media for 30 min. For stress granule analogues, cells were transfected as described above with RRM-CR_x5_ or RRM-CR_x5_ (pMT2) using cytoYF_x5_. The cells were allowed to grow for 36-48 h before rapamycin administration. The rest of the experiment was carried out as described above.

### Electron microscopy

For correlative EM, cells were cultured on sapphire disks that are carbon-coated with a grid pattern. This pattern was used to locate the region of interest in an electron microscope. Following poly-D-lysine coating, cells were cultured overnight on the sapphire disks in a 24-well culture plate before the transfection. The iPOLYMER formation was induced 24 h after transfection for 1 h at 333 nM rapamycin, and cells were fixed with 4% paraformaldehyde (in PBS) for 5 minutes. Cells were subsequently washed with PBS three times and then imaged on a fluorescence microscope in the culture plate. Fluorescence images of iPOLYMER puncta-containing cells were obtained with a 20x objective lens. Bright field images of the carbon grid pattern on the sapphire disks were simultaneously obtained to locate the cells in later steps.

Following fluorescence imaging, cells were prepared for EM. To avoid shrinkage in cells during the dehydration process, cells were frozen on a high-pressure freezer (Leica EM ICE). The vitrified specimens were handled carefully under liquid nitrogen and placed into cryo-tubes containing 1% osmium tetroxide (EMS #19132), 1% glutaraldehyde (EMS #16530), 1% water, and anhydrous acetone (EMS #10016). The freeze-substitution was performed in a Leica AFS2 unit with the following program: 8 h at −90^<u>o</u>^C, 5^<u>o</u>^C/h to −20^<u>o</u>^C, 12 h at −20^<u>o</u>^C, and 10^<u>o</u>^C/h to 20^<u>o</u>^C. The specimens were then embedded into epon-araldite resin and cured for 48 h. The region of interest was located based on the grid pattern. The plastic block was trimmed to the region of interest and sectioned on a diamond knife using an ultramicrotome (Leica UC8). Approximately 40 consecutive sections (40 nm each) were collected onto the pioloform-coated grids and imaged on a Phillips transmission EM (CM120) equipped with a digital camera (AMT XR80). The fluorescence and electron micrographs were roughly aligned based on the carbon-coated grid patterns. The alignment was slightly adjusted based on the visible morphological features. EM images of the stress granules were obtained similarly, but without any correlation.

### Immuno-staining

Cells were fixed with 4% (w/v) paraformaldehyde, permeabilized with 0.2% (v/v) Triton X-100 for 30 min, and blocked by 1% BSA in PBS. A monoclonal antibody against PABP-1 (10E10; Sigma-Aldrich), G3BP1 (sc-81940; Santa Cruz), eIF4G (sc-11373; Santa Cruz), and eIF3b (sc-16377; Santa Cruz) were used at a 1:500-800 dilution. The secondary antibodies were Alexa Fluor 594- or 647- labeled anti IgG of the appropriate species (anti-human antibodies were purchased from Jackson Immunoresearch, others from Invitrogen), with a 1:1000 dilution. For longer administration conditions (24 h), rapamycin was used at 100 nM. After the staining, cells were imaged with either a Zeiss Axiovert 135 TV microscope equipped with a QIclick camera (QImaging), or a Nikon eclipse Ti microscope equipped with a Zyla sCMOS camera (Andor). For imaging Alexa Fluor 594 and 647, filter sets for mCherry and Cy5 were used, respectively.

### Light-inducible iPOLYMER-LI puncta formation

For live-cell imaging, cells were transfected with mCherry-iLID_x6_ and YFP-SSBP_x6_, incubated for 12-24 h, then observed under the LSM780 confocal microscope (Zeiss). Using the bleaching function of the microscope, cells were stimulated by a 488 nm laser before each frame during the stimulus. For imaging, YFP and mCherry signals were excited by 514 nm and 564 nm lasers, respectively.

For immunostaining study of iPOLYMER-LI puncta, we used a custom-made blue light LED illuminator for stimulation. Cells were transfected with mCherry-iLID_x6_ and YFP-SSBP_x6_ or the constructs functionalized with TIA-1 RRM domains, incubated for 24 h, and stimulated by the illuminator in the incubator for 1 h. Cells were then fixed and immunostained as described above.

### Statistical analysis

A two-tailed Student's *t*-test was used for all statistical analyses. In figures, S.E.M. was used to generate error bars unless the deviation was too small to be visible on the plot.

## Acknowledgements

We are grateful to J. L. Pfaltz for kindly working with A.S.A. to develop a modified C++ code for identifying chordless cycles in graphs. We are thankful to N. Kedersha and P. Anderson for helpful discussions and reagents related to stress granules. We appreciate R. Reed, A. Ewald, H. Sesaki, M. Iijima and S. Regot for sharing their resources for the experiments. This work was supported by the National Institutes of Health (NIH) (GM092930, DK102910, CA103175 and DK089502 to T.I., and T32GM007445 to A.S.), the National Science Foundation (NSF) (CCF-1217213 to J.G.), and the Johns Hopkins University Catalyst Fund to T.I..

## Author Contributions

H.N., A.L. and T.I. conceived the project. H.N., A.L., A.S., Y.C.L., M.T., R.D., D.B., performed molecular biology as well as cell biology experiments. H.N., A.L., S.R. and A.S. purified proteins under guidance by W.H. and S.B.G.. The biochemical and biophysical experiments were mostly performed by H.N and A.L., and partially by S.R. and Y.C.L.. H.N., A.L., and T.I. wrote the manuscript with the help of J. G.. A.S.A and J.G. developed the computational model, analyzed the computational results, and wrote the computational parts of the paper. A.S.A. wrote appropriate code and conducted the computational experiments. S.W. performed correlated EM measurement and analysis. E.R. and B. H. performed development and demonstration of light-inducible iPOLYMER with H. N.

## Competing Financial Interests statement

The authors declare no financial interest associated with the present work.

